# Efficacy of antiviral drugs against the omicron variant of SARS-CoV-2

**DOI:** 10.1101/2021.12.21.473268

**Authors:** Agnieszka Dabrowska, Artur Szczepanski, Paweł Botwina, Natalia Mazur-Panasiuk, Helena Jiřincová, Lukasz Rabalski, Tomas Zajic, Grzegorz Popowicz, Krzysztof Pyrc

## Abstract

The Omicron variant of the SARS-CoV-2 virus was first detected in South Africa in November 2021. The analysis of the sequence data in the context of earlier variants suggested that it may show very different characteristics, including immune evasion and increased transmission. These assumptions were partially confirmed, and the reduction in protection in convalescent patients and vaccinated individuals have been confirmed. Here, we have evaluated the efficacy of antivirals against SARS-CoV-2 variants, Omicron, Delta, and the early 2020 isolate.

## Text

The SARS-CoV-2 virus emerged in 2019 in South-Eastern Asia, to spread rapidly in 2020 to all continents and cause the global pandemic. In 2020, several variants of the virus emerged, but none became dominant for a long time; however, the end of 2020 brought us the alpha variant that emerged in the United Kingdom, to cause the winter wave of infections in many countries. However, its prime position was abolished during the spring/summer season by the delta variant, first detected in India in the late 2020. These viruses were characterized mainly by the increased transmissibility, as the need for more effective transmission mainly drove their evolution. In the meantime, several variants evading the immune responses were identified. The beta variant was first detected in autumn 2020, Lambda was in August 2020, and Mu in January 2021. All these variants carried mutations localized to the epitopes recognized by the neutralizing antibodies. However, due to evolutionary inferiority, they have never become dominant^1^. In November 2021, the world was electrified by the information on the emergence of a new variant, which carried mutation previously associated with evasion of the immune responses. At the same time, some suggested also improved transmissibility. The epidemiological information on the rapid spread of the virus in South Africa and first reports showing its decreased susceptibility to the antibody-mediated neutralization strongly reinforce these theses. Therefore, Omicron (B.1.1.529) variant poses a threat to public health, leading to serious suppression in global pandemic control^2, 3^. In this study, we evaluated antivirals’ efficacy against selected SARS-CoV-2 variants, Omicron, Delta, and the isolate from May 2020.

First, we verified the identity of isolates after the recovery in cell culture. The virus isolated *in house* in spring 2020 in Poland was used as a reference (B.1.13; designated hCoV-19/Poland/PL_P7/2020; GISAID accession code: EPI_ISL_428930). The Delta variant (B.1.617.2) was isolated from a sample obtained in May 2021 in the Czech Republic (hCoV-19/Czech Republic/NRL_7102/2021; GISAID accession code: EPI_ISL_2357738). The Omicron variant [B.1.1.529; BA.1] was isolated from a sample obtained in December 2021 in the Czech Republic [hCoV-19/Czech_Republic/KNL_2021-110119140/2021; GISAID accession code: EPI_ISL_6862005]. Omicron and Delta variants were originally isolated on Vero E6 cells overexpressing TMPRSS2 (NIBSC: 100987). Viral stocks were generated by infecting monolayers of Vero cells (ATCC CCL-81). Virus yields were assessed by titration according to the method of Reed and Muench.

The Omicron and Delta variants were propagated for 2 and 5 passages, respectively, and subjected to whole-genome sequencing to confirm genomic sequence identity to the original isolates. Total RNA was extracted from cell culture supernatants using MagnifiQ™ Viral RNA kit (AA Biotechnology, Gdańsk, Poland) and automated nucleic acid isolation system KingFisher Flex 96 deep well plate (Thermo Fisher Scientific, Waltham, MA, USA) according to the manufacturer’s guidelines. Whole-genome sequencing was performed using Oxford Nanopore Technique Technologies (ONT) platform (Oxford, United Kingdom) with the ARTICv1200 sequencing protocol. Obtained sequences are deposited at the GISAID database under the following accession names: hCoV-19/Czech Republic/KNL_2021-110119140/2021_p2 for Omicron hCoV-19/Czech Republic/NRL_7102/2021_p5 for Delta. No significant differences in sequence were recorded in the hotspots described for these lineages.

As it is known that the efficacy of vaccines and some monoclonal antibodies is reduced, it is of importance to verify whether the small molecule drugs targeting the more conserved proteins remain effective. We focused mainly on the drugs that went successfully through the clinical trials – Legevrio by Merck (polymerase inhibitor; molnupiravir^4^, EIDD-2801; MK-4482, MedChemExpress HY-135853 and its derivative beta-d-N4-hydroxycytidine^5^, MedChemExpress HY-125033) and Paxlovid by Pfizer (protease inhibitor; PF-07321332^6^; MedChemExpress HY-138687). We have also included acriflavine^7^ (SigmaAldrich; A8126), which we previously shown to be a potent *ex vivo* and *in vivo* inhibitor of betacoronaviruses blocking the PL^pro^ protease. Further, remdesivir^8^ sold by Gilead Sciences as Veklury by was included in the panel (polymerase inhibitor; GS-441524; Cayman Chemical, USA) and AT-527 developed by Atea Pharmaceuticals^9^ (polymerase inhibitor; bemnifosbuvir hemisulfate; MedChemExpress HY-137958). All inhibitors were dissolved in DMSO (Sigma-Aldrich, Poland) to the final concentration of 10 mM.

To test the antiviral activity of the compounds, subconfluent A549^ACE2+TMPRSS2+^ cells^10^ were infected with viruses (all three variants) at 1600 50% tissue culture infectious dose (TCID50)/ml in the presence of selected concentrations of inhibitors. Control was prepared in the same manner, but no inhibitor was added. After 2 h of incubation at 37°C, cells were rinsed twice with PBS, and a fresh medium with compounds was added. The infection was carried out for 72 h, and the cytopathic effect (CPE) was assessed. Culture supernatants were collected for analysis, which was carried out as previously described^7^. The experiment was performed in three biological repetitions, each in duplicate (n=6). The half maximal inhibitory concentration (IC_50_) value was calculated using GraphPad Prism 9.0. Obtained results are presented in Figure 1A-F and the estimated IC_50_ values are presented in Figure 1G. For paxlovid, molnupiravir, acriflavine and remdesivir the observed inhibition and IC_50_ was similar as described previously, and maintained for all the variants. We did not, however, observe the activity of AT-527, but this is most likely associated with the model itself and the inability of cells to process the inert pro-drug to its active metabolite. Concluding, the obtained results show that the drugs that are being developed against the SARS-CoV-2 will likely retain its efficacy also for the omicron variant.

**Figure 1.**
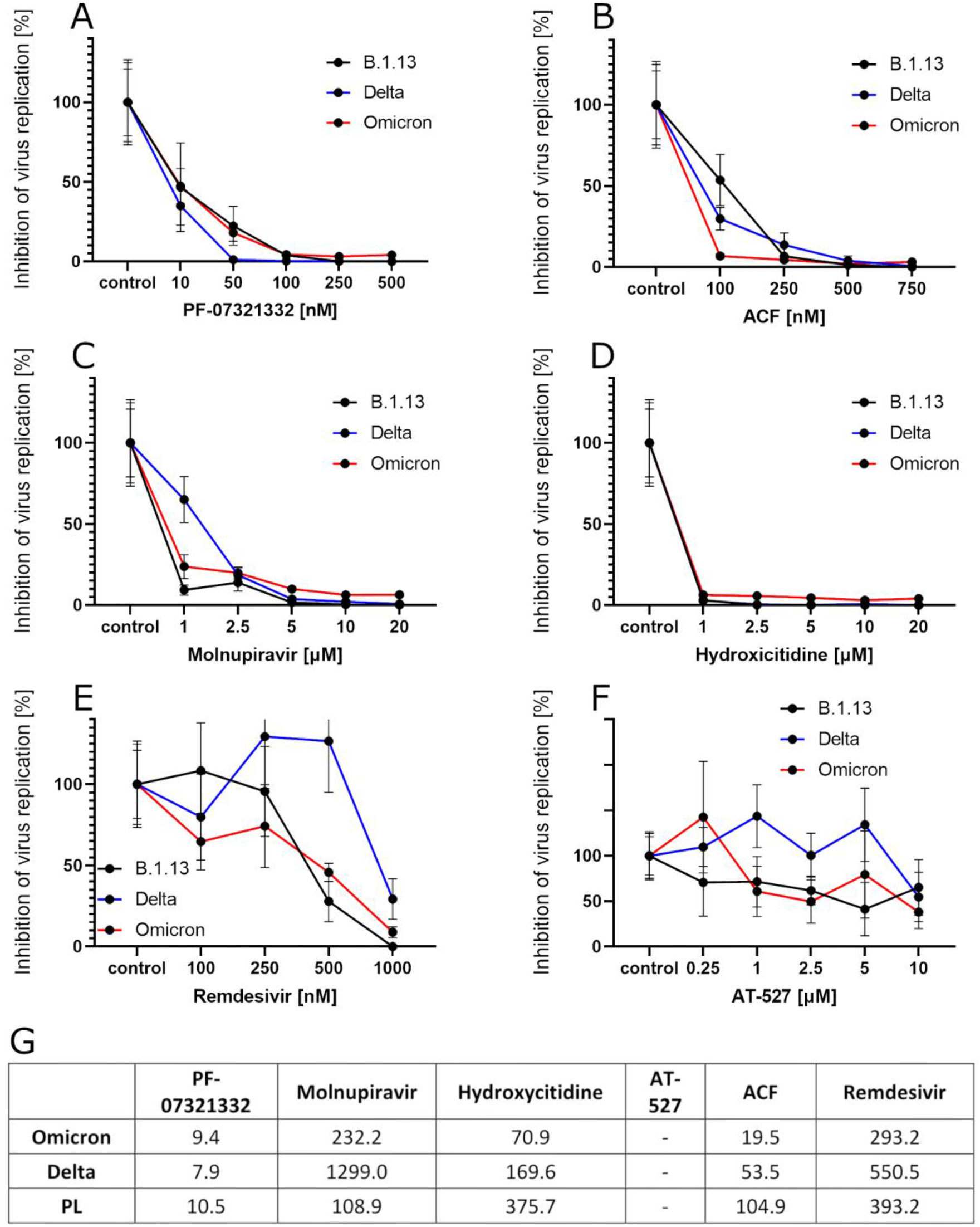
Efficacy of SARS-CoV-2 inhibitors against the omicron variant. **(A)-(F).** Dose dependent inhibition of the virus replication, as measured by the RT-qPCR. Virus yield was normalized to the control samples and the inhibition at given concentration is given as percent. **(G)** IC_50_ values of the tested compounds.

## Acknowledgments

This work was supported by the subsidy from the Polish Ministry of Science and Higher Education for the research on the SARS-CoV-2, a grant from the National Science Center UMO-2017/27/B/NZ6/02488, and the Corona Accelerated R&D in Europe (CARE) project. The CARE project has received funding from the Innovative Medicines Initiative 2 Joint Undertaking (JU) under grant agreement No 101005077.

